# XRN1-dependent RNA homeostasis regulates hepatitis D virus

**DOI:** 10.64898/2026.11.01.763535

**Authors:** Senko Tsukuda, Medhi Boutasbih, Peter A.C. Wing, Aleksandr Fedorov, Xiaxuan Zhu, Jan Rehwinkel, Peter Balfe, Jane A McKeating

**Affiliations:** Chinese Academy of Medical Sciences Oxford Institute, University of Oxford, UK; MRC Translational Immune Discovery Unit, MRC Weatherall Institute of Molecular Medicine, Radcliffe Department of Medicine, University of Oxford, Oxford, UK; Nuffield Department of Medicine, University of Oxford, Oxford, UK

**Author notes:** Department of Virology II, National Institute of Infectious diseases, Japan Institute for Health Security, Tokyo, Japan. Shared communicating authors (Senko Tsukuda, Jane A McKeating).

## Abstract

Hepatitis delta virus (HDV) infection causes the most severe form of viral hepatitis, yet the host mechanisms restricting HDV replication remain poorly understood. Although HDV infection activates the MDA5-LGP2 pathway and induces robust interferon (IFN) responses, these antiviral programs have limited impact on viral replication. Here we identify the 5’-3’ exoribonuclease XRN1 as a key regulator of HDV infection. Loss of XRN1 in hepatoma cells increased double-stranded RNA (dsRNA) levels and induced interferon-stimulated genes (ISGs), including MDA5 and MAVS, and limited HDV infection. XRN1 deficiency enhanced ADAR1-mediated RNA editing, revealing a dynamic interplay between endogenous dsRNA metabolism and antiviral sensing. Inhibition of type I IFN signaling restored MDA5 expression and rescued HDV replication in XRN1-deficient cells. To identify the downstream effectors responsible for this restriction, we combined transcriptomic profiling with functional siRNA screening and identified DUSP5 and SIRPA as previously unrecognized ISGs with anti-HDV activity. Together, our findings establish XRN1 as a gatekeeper of dsRNA homeostasis that controls HDV replication through innate immune activation and reveals host antiviral effectors that may provide therapeutic opportunities for chronic hepatitis D.

**Graphical abstract:** Activation of dsRNA sensing pathway following XRN1 depletion upregulates IFN production and IFN downstream pathways, leading to increases in host defenses to HDV infection. In addition XRN1 depletion increases ADAR1 expression, which increases L-HDAg levels and suppresses HDV replication. Image created in BioRender : https://BioRender.com/t73u45i

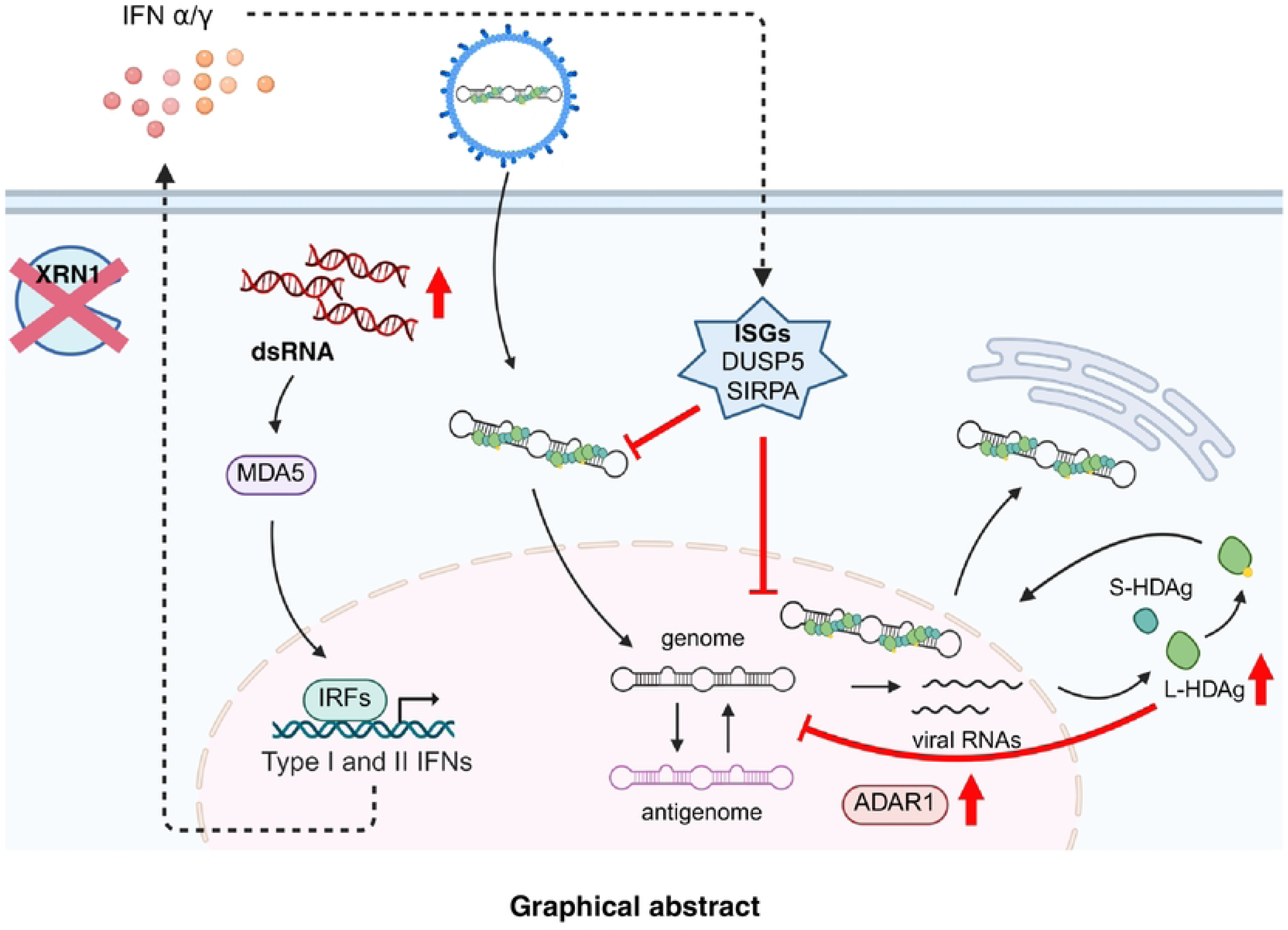

**AUTHOR SUMMARY:** Hepatitis D virus causes the most severe form of viral hepatitis, yet how our cells defend against this virus remains poorly understood. Although infection triggers strong immune responses, their effects on viral replication are limited. In this study, we identified XRN1, an enzyme involved in cellular RNA degradation, as an important regulator of hepatitis D virus infection. We found that loss of XRN1 causes the accumulation of double-stranded RNA, activates antiviral immune responses, and suppresses viral replication. We also observed increased RNA editing, highlighting the connection between RNA metabolism and antiviral immunity. Furthermore, we identified two previously unrecognized antiviral factors, DUSP5 and SIRPA, that contribute to controlling viral infection. Our findings reveal how cellular RNA degradation regulates antiviral defenses and suggest potential new approaches to treating chronic hepatitis D.

## INTRODUCTION

Hepatitis D virus (HDV) is a major health problem with >12 million cases worldwide and is associated with an accelerated course of liver disease. Approximately 5% of chronic hepatitis B virus (HBV) infected individuals are co- or super-infected with HDV^1,2^. HDV encodes a small (1.7kb) circular, single-stranded, negative-sense RNA that replicates via a double rolling circle mechanism and requires HBV envelope proteins (HBs) to assemble infectious particles. HBV vaccines prevent HBV and HDV co-infection, but there are no HDV specific vaccines to prevent super infection^2,3^. The immunomodulator pegylated IFN-α (Peg-IFN-α) was the only recommended therapy for chronic hepatitis D prior to the European and the U.S. Food and Drug Administration (FDA) approval of the entry inhibitor Bulevirtide^3–5^. However, Peg-IFN-α is only effective in 20-30% of cases and is frequently associated with significant side effects and viral rebound post therapy^6,7^.

Discrimination between self and non-self RNA is essential for effective antiviral immunity and to limit aberrant inflammation and autoimmunity. During viral infection, cytoplasmic double-stranded RNAs (dsRNAs) are sensed by the pattern-recognition receptors retinoic acid-inducible gene I (RIG-I), melanoma differentiation-associated protein 5 (MDA5), and laboratory of genetics and physiology 2 (LGP2), leading to the activation of the adaptor mitochondrial antiviral signaling protein (MAVS). MAVS signaling induces IFN production and the expression of interferon-stimulated genes (ISGs) to induce an antiviral state^8,9^.

Recent studies have shown that epigenetic changes in cancer cells are associated with an accumulation of dsRNA and activated antiviral signaling in the absence of infection, a process termed ‘viral mimicry’. Disrupting epigenetic regulation^10–13^, RNA splicing^14^ or protein arginine methylation^15^ promotes the transcription of transposable elements (TE) that activate dsRNA sensing pathways and lead to cancer cell death. The 5′–3′ exoribonuclease 1 (XRN1) degrades both single-stranded RNA (ssRNA) and dsRNA bearing a 5′ monophosphate and has a central role in cellular RNA turnover^16,17^. XRN1 has also been reported to degrade viral RNAs and contribute to host antiviral defenses^18–21^.

HDV infection is sensed by MDA5 in cooperation with LGP2 triggering the expression of IFN-β, IFN-λ and a broad range of ISGs^22,23^. Importantly, these antiviral responses have a modest effect on HDV replication^23,24^, and the ISGs responsible for restricting HDV are poorly defined^25^. In this study, we demonstrated that ISGs induced following XRN1 depletion restrict HDV replication. Transcriptomic analysis, combined with a small interfering RNA (siRNA) screen, identified dual specificity phosphatase 5 (DUSP5) and signal-regulatory protein alpha (SIRPA) as anti-HDV ISGs. These findings provide new insights into host antiviral responses against HDV and suggest potential new therapeutic strategies.

## RESULTS

### Depletion of XRN1 in HepG2 hepatoma cells activates IFN signalling

Genetic disruption of XRN1 in some cancer cell lines increased dsRNA levels and activated the dsRNA sensors MDA5 and protein kinase R (PKR)^26,27^ (**Fig.1B**). To assess the role of XRN1 in human hepatoma cells we evaluated dsRNA levels in parental (wild type, WT) and knockout (KO) cell lines (KO22 and KO35)^28^ using a dsRNA specific antibody. Both KO lines showed elevated dsRNA levels relative to the WT cells together with an upregulation in the RNA sensors, RIG-I and MDA5, and their adaptor protein MAVS: HepG2-NTCP WT and KO cells (**Fig.1A**). To detect IFN expression, we used the reporter HEK293 cell line expressing firefly luciferase under the control of an interferon-stimulated response element^29^, and noted increased activity following treatment with the supernatant from both KO cell lines (**Fig.1A**). Both KO cells showed increased expression of STAT1 and phosphorylated-STAT1 (pSTAT1) (**Fig.1A**), consistent with IFN induction. PKR activation in XRN1 depleted lung adenocarcinoma A549, non-small cell lung carcinoma NCI-H1229, NCI-H460, and NCI-H1437 cell lines has been shown to increase IFN-β induced cellular lethality^27^. Both HepG2 XRN1 KO cells showed increased levels of PKR and its activated form, phosphorylated PKR (pPKR). We investigated the sensitivity of the KO cells to IFN-α, IFN-β and IFN-γ and showed an increase in cytotoxicity relative to the WT cells (**Fig.1B**). Collectively, these data show that dsRNA upregulation in XRN1 KO enhances both MDA5- and PKR-signalling pathways. Inspecting the transcriptome of HepG2 KO22 cells identified significant up-regulation of 123 genes from a curated list of ISGs^30,31^ (**Fig.1C**). Gene set enrichment analysis (GSEA) of these genes showed an upregulation of pathways associated with IFN-α and IFN-γ responses (**Fig.1D, Supplementary Table 1**).

**Figure. 1.**
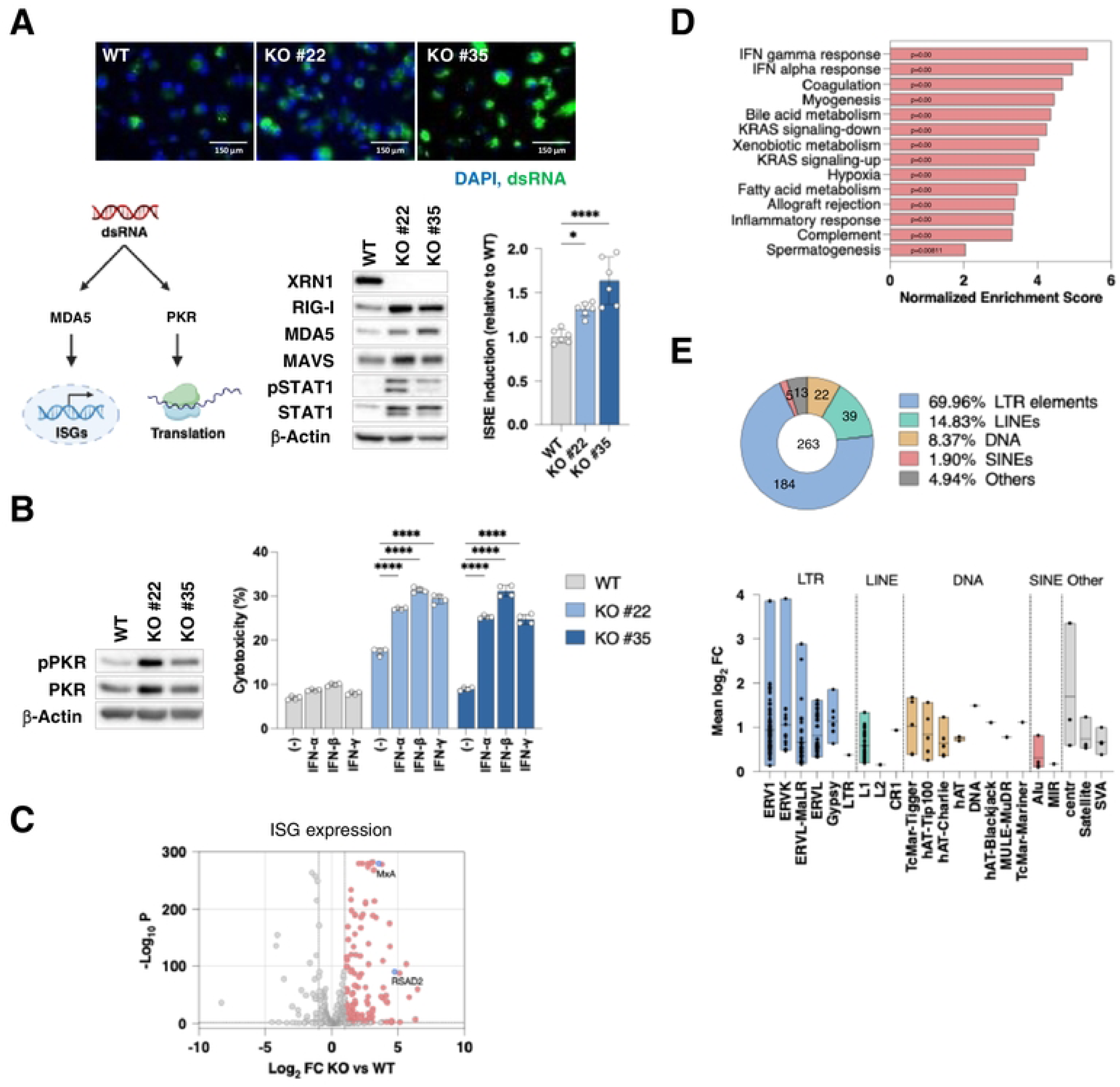
XRN1 depletion induces intracellular dsRNA sensing. (**A**) **Detection of dsRNA in XRN1 KO cells.** XRN1 HepG2-NTCP WT and XRN1 knockout clones #22 and #35 (KO #22 and KO #35) cells were cultured for 9 days under differentiation conditions. Double-strand RNA (dsRNA) (green) and nuclear DNA (blue) were detected by immunofluorescence staining (scale bar = 150 µm). Protein levels of XRN1, RIG-I, MDA5, MAVS, pSTAT1, STAT1 and β-Actin were assessed by western blotting. HEK293T cells stably expressing an IFN-stimulated response element (ISRE)-derived reporter were incubated with cell culture supernatant from HepG2-NTCP XRN1 WT or KO cells for 48h and reporter activity evaluated by a luciferase assay. Data from 6 independent samples are shown as mean ± SD and analyzed using Mann-Whitney tests with Bonferroni correction. (**B**) **PKR-mediated IFN sensitivity in XRN1 KO cells.** HepG2-NTCP cells were differentiated for 6 days then treated with 1000 IU/mL of IFN-α, 100 ng/mL IFN-β, or 100 ng/mL IFN-γ for 3 days. Protein levels of pPKR, PKR and β-Actin were assessed prior to IFN treatment by western blotting. Cytotoxicity was evaluated by an LDH assay. Data is expressed relative to a positive control treated with 0.05% Triton-X100 (100% cytotoxicity). (mean ± SD, n = 9 from 3 independent experiments). **** *p* < 0.0001. (**C-D**) **ISG expression and upregulated pathways in XRN1 KO cells.** HepG2-NTCP WT and XRN1 KO #22 cells were cultured for 9 days under differentiation conditions, RNA was extracted, DNase treated and analyzed by RNAseq. (**C**) **ISG expression by volcano plot.** The X and Y axes show the log_2_ fold change (FC) and -log10 (p-value) of the changes, respectively. The dotted lines show the thresholds, defined as log_2_ fold = ±1 and -log_10_[p] = 1.303. MxA and RSAD2 are indicated by blue dots. (**D**) **GSEA hallmark analysis.** RNA-seq of WT and XRN1 KO22 cells identified upregulated pathways in the KO #22 cells, the normalized enrichment scores are shown. (**E**) **Upregulated transcripts from transposable elements in KO #22 cells. The** percentage of each transposable element (TE) class, including long terminal repeats (LTRs), long interspersed elements (LINEs), DNA derived TE transcripts, and short interspersed nuclear elements (SINEs) that were significantly upregulated in KO cells relative to WT cells (P value < 0.05) are shown, with the numbers of identified TE transcripts indicated. The mean log_2_FC in expression of members of each TE class, the center line and box limits indicate the median, maximum and minimum Mean log_2_FC, respectively.

Transposable elements (TEs), including long terminal repeats (LTRs), long interspersed elements (LINEs), and short interspersed nuclear elements (SINEs), constitute a substantial proportion of human genome and represent an important source of endogenous RNAs capable of forming immunostimulatory dsRNA^27,32–36^. Aberrant expression or accumulation of TE-derived RNAs can activate cytoplasmic dsRNA sensors and trigger innate immune responses. However, the impact of XRN1 depletion on TE gene expression in hepatoma cells is not known. To address this, we mapped TE transcript expression in WT and KO22 cells (BioProject ID: PRJNA1228332). 263 TE subfamilies were significantly upregulated in KO22 cells, including LTRs (70 %), LINEs (15%), DNA derived (8 %) and SINEs (2%). We also noted a significant increase in endogenous retroviral gene expression in KO22 cells (**Fig.1E**). These data support a model where dsRNAs derived from accumulated TE-containing transcripts activate RNA sensors and down-stream IFN signalling in XRN1 depleted cells.

### A differential role for XRN1 in the regulation of HDV and other RNA viruses

To assess a role for XRN1 in HDV replication we infected WT and XRN1 KO cells with HDV and quantified viral RNA, viral antigenome and HDAg at 6 days post infection (dpi). We observed a significant reduction in both HDV RNA and the frequency of HDAg expressing cells in the XRN1 KO cells. Although several ISGs known to be induced by HDV infection, including myxovirus resistance protein A (MxA) and radical S-adenosyl methionine domain-containing 2 (RSAD2)^22,23^, were constitutively upregulated in XRN1 KO cells (**Fig. 1C**, blue dots), their further induction following HDV infection was attenuated, consistent with the reduced levels of HDV replication (**Fig.2A**). As our quantitative PCR (qPCR) cannot discriminate between viral antigenome and mRNA we assessed HDV antigenomes by single molecule inexpensive fluorescence *in situ* hybridization (smiFISH) using a published protocol^37^. Co-staining of HDAg and antigenome revealed clear co-localization of both signals with a significant reduction of antigenome levels in the XRN1 KO22 cells (**Fig.2A**). Importantly, in both KO cells the antiviral phenotype was reversed by exogenous expression of XRN1 (**Fig 2B**).

**Figure. 2.**
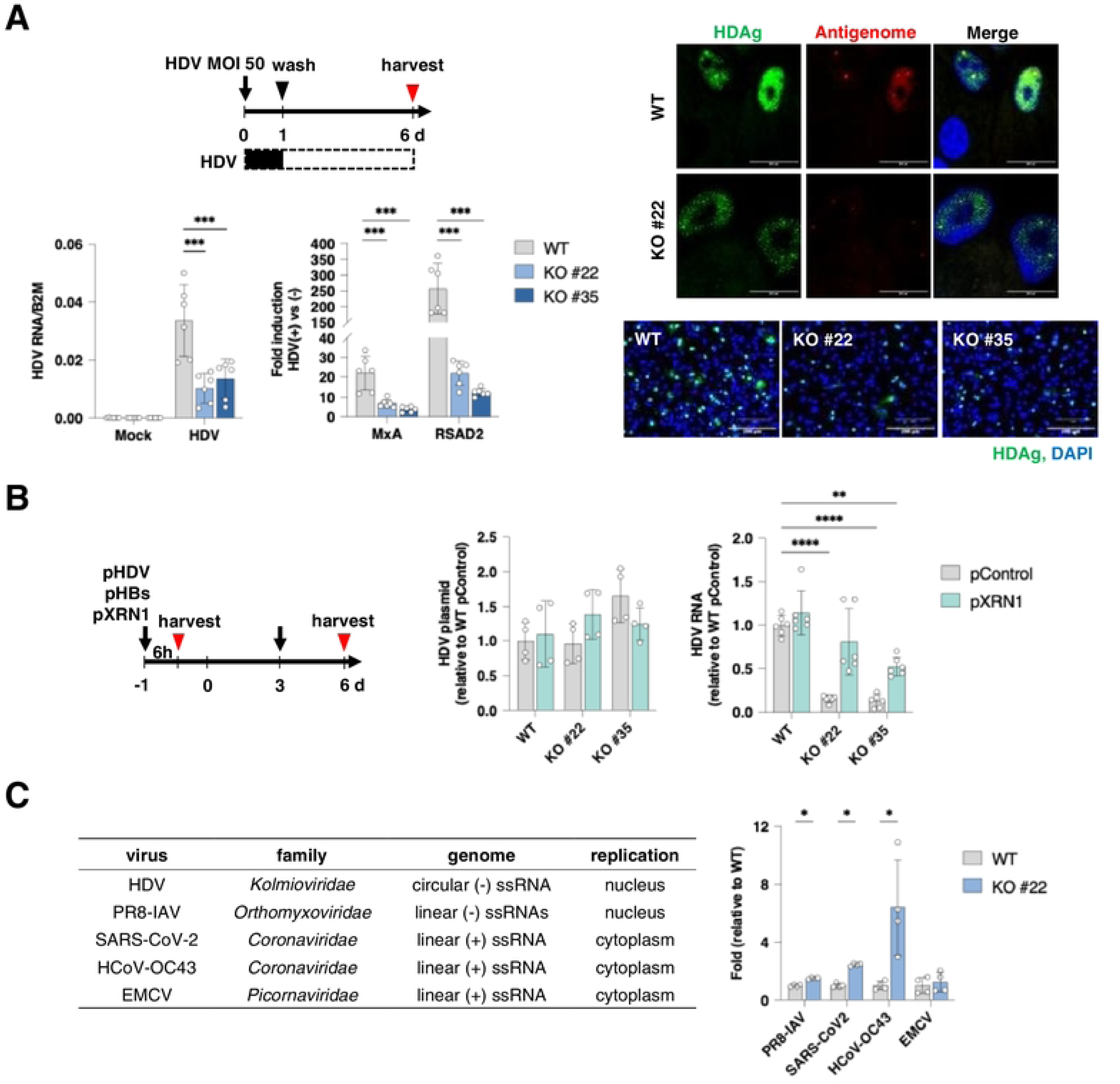
Disruption of XRN1 attenuates HDV replication. (**A**) **HDV replication in XRN1 WT and KO cells.** HepG2-NTCP WT and KO cells were differentiated for 3 days and then infected with HDV (MOI = 50). After 24h, the cells were washed and incubated for 6 dpi. RNA was extracted from the cells and HDV RNA detected by RT-qPCR using primers targeting the antigenome or mRNA. HDV RNA levels were normalized to a housekeeping gene, *B2M*. The levels of *MxA* and *RSAD2* induced by HDV infection in each cell line were normalized in the same way (mean ± SD, n=6 from 3 independent experiments). HDAg (Green) and nuclei (Blue) were detected by immunofluorescence staining, one image set from 3 independent experiments are shown. The HDV antigenome was detected by single molecule inexpensive FISH (smiFISH) using previously published probes^37^. HDAg (Green), HDV antigenome (Red) and nuclei (Blue) are shown, scale bar = 20 µm. The images were obtained from one of 2 independent infection experiments. (**B**) **HDV replication in XRN1 over-expressing cells.** HepG2-NTCP WT and KO cells were transfected with plasmids encoding HDV (pHDV) and Hepatitis B virus surface protein (pHBs), together with XRN1 (pXRN1) or a control plasmid (pControl) for 6h, medium was changed and cells cultured for 6 days. To maintain expression levels, pXRN1 and pControl were also transfected at 3 days. DNA was extracted at 6h post-transfection and HDV plasmid levels quantified by qPCR with normalization to a housekeeping gene, *PrP* (mean ± SD, n = 4 from 2 independent experiments). At 6 days post-transfection, RNA was extracted, TURBO DNase treated and RNA levels assessed by RT-qPCR with normalization to the housekeeping gene, *B2M* (mean ± SD, n = 6 from 3 independent experiments, Mann-Whitney tests with Bonferroni correction * *p* < 0.05, ** *p* < 0.01, \*\*\**p* < 0.005, **** *p* < 0.0001). (**C**) **Effect of XRN1 KO on RNA viruses.** HepG2-NTCP cells were seeded on collagen coated plasticware or collagen-coated plates and after 3 days infected with Influenza A virus A (IAV) PR8-GFP, severe acute respiratory syndrome coronavirus 2 (SARS-CoV-2) (VIC-01 mNeonGreen), human coronavirus OC43 (HCoV-OC43) or Encephalomyocarditis virus (EMCV) at an MOI of 1. PR8-GFP and SARS-CoV-2 infections were quantified at 24h post infection by measuring GFP fluorescence and normalised by DAPI counter staining. For OC43 and EMCV viral titres were quantified by plaque assay on Vero cells.

To assess whether other single-strand RNA (ssRNA) viruses were sensitive to XRN1 depletion we infected WT or XRN1 KO22 cells with a range of viruses that replicate in the nucleus or cytoplasm, including: influenza A virus (PR8-IAV, *Orthomyxoviridae*); severe acute respiratory syndrome coronavirus 2 (SARS-CoV-2, *Coronaviridae*); human coronavirus OC43 (HCoV-OC43, *Coronaviridae*) and encephalomyocarditis virus (EMCV, *Picornaviridae*). XRN1 depletion had a minimal effect on the replication of the other viruses tested (**Fig 2C**). Our data shows that XRN1-dependent dsRNA sensing in hepatoma cells selectively restricts HDV replication and challenges the concept of generalised ‘viral mimicry’.

### XRN1 depletion increases RNA editing by ADAR1

RNA editing by adenosine deaminase acting on RNA 1 (ADAR1) is an essential step in the HDV life cycle and edits the amber stop codon of the HDV antigenome, converting a UAG codon (Stop) to UGG (Tryptophan). This conversion enables expression of the large HDAg isoform (L-HDAg), essential for virion assembly and is reported to suppress viral RNA replication (**Fig.3A**). ADAR1 encodes two isoforms, p110 (110 kDa) and p150 (150 kDa), and XRN1 depletion leads to a relative increase in p150, consistent with the presence of an IFN-inducible promoter driving p150 expression (**Fig.3B**). Infecting the WT or KO22 cells with increasing multiplicities of infection (MOI) showed a dose-dependent reduction in L- and S-HDAg expression in the KO cells and an increase in the relative amount of L-HDAg (**Fig.3B**). To assess whether ADAR1 editing was increased in XRN1 depleted cells, we analysed viral and host mRNA editing sequences in our poly(A) RNA-seq data. Consistent with the HDAg protein levels, XRN1 KO22 cells showed higher levels of U**A**G stop to U**G**G tryptophan codon conversion in HDAg encoding mRNA (**Fig.3C**). Importantly, XRN1 depletion altered A-to-I editing of host transcripts in uninfected cells. Total RNA-seq revealed that most ADAR1editing sites detected in both the WT and KO cells, primarily in 3′ untranslated regions (UTRs), showed significantly increased editing (964 sites), whereas only 17 sites showed a reduction. A similar pattern was observed for sites detected in only one cell type: 5,680 sites were specific to XRN1-depleted cells and 1,815 to parental cells (**Fig.3D**). In conclusion, XRN1 depletion increased ADAR1 expression and perturbed HDV and host RNA editing.

**Figure. 3.**
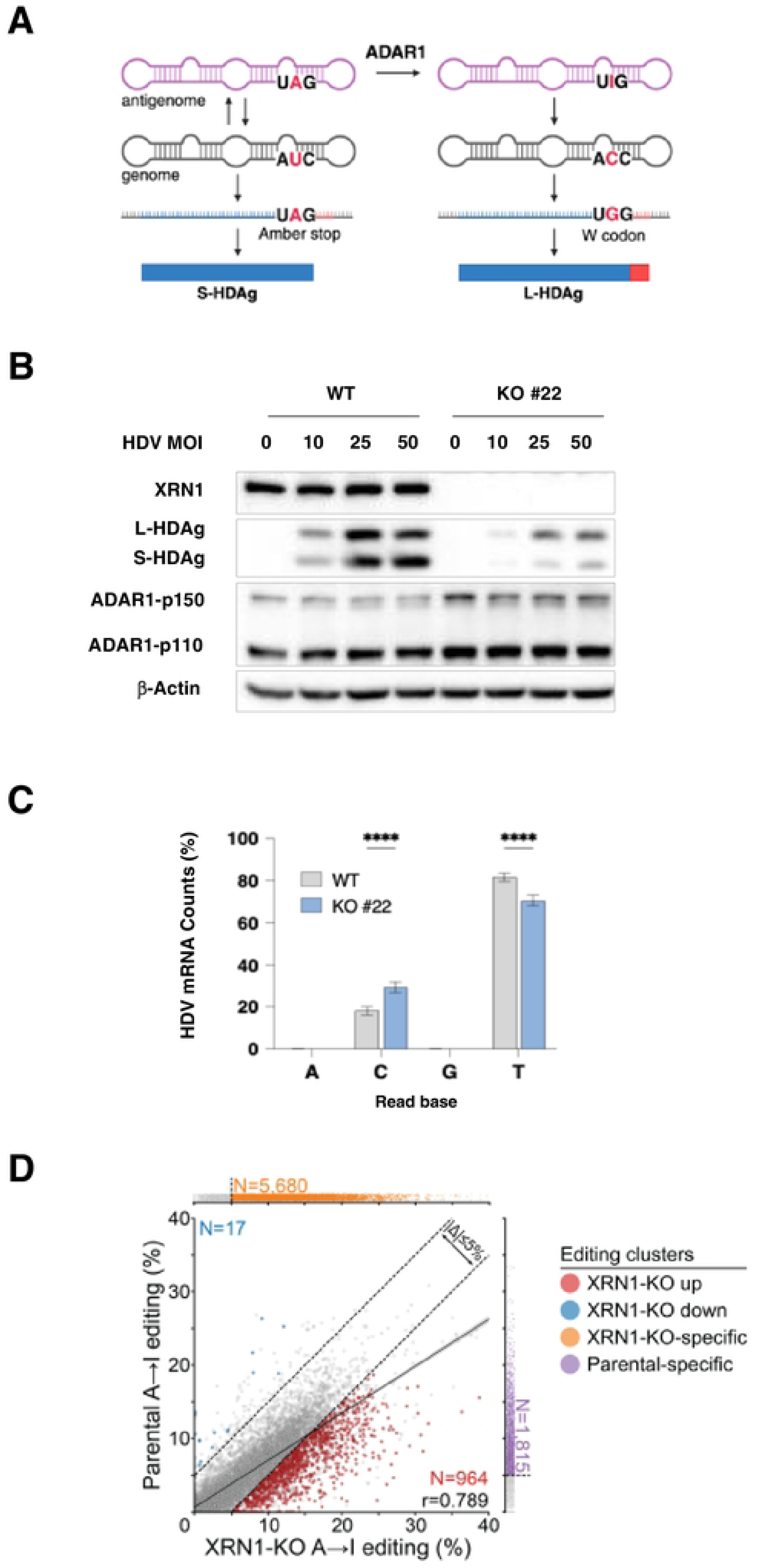
XRN1 regulates ADAR1 expression and function. (**A**) **Schematic of HDV RNA-editing by ADAR1.** (**B**) **Protein levels of ADAR1 and HDAg isoforms in HepG2-NTCP WT and KO cells.** Cells were differentiated for 3 days and infected with HDV (MOI of 0, 10, 25 or 50). After 24 h the cells were washed and incubated for 6 days. XRN1, L-HDAg, S-HDAg, ADAR1-p150, ADAR1-p110 and β-Actin levels were detected by western blotting. (**C**) **RNA editing by ADAR1 of viral and host mRNAs.** Differentiated HepG2-NTCP WT or XRN1 KO cells were inoculated with HDV (MOI = 50) for 24 h, washed, and cultured for up to 6 dpi under differentiation conditions. RNA was extracted, treated DNase and analyzed by RNA-seq. **(D) Effect of XRN1 loss on A-to-I editing of host RNAs**. Comparison of median A-to-I editing frequencies between uninfected HepG2-NTCP WT and KO #22 cells, where each point represents a cluster. The main panel shows clusters detected in both conditions, with statistically significant differences highlighted in color. The marginal panels show clusters detected in only one condition. The solid line shows the linear regression fit and shading indicates the 95% confidence interval, *r* is the Pearson correlation coefficient. Dashed lines indicate differences of 5% between conditions in the main panel and editing frequencies of 5% in the marginal panels.

### Inhibition of type I IFN signalling restores MDA5 expression and HDV replication in XRN1 depleted cells

We hypothesized that the increased dsRNA levels and associated IFN signalling suppresses HDV replication. To assess this, cells were treated with the type I IFN inhibitor, B18R, and evaluated for RIG-I and MDA5 expression together with the downstream JAK-STAT pathway (**Fig.4A**). B18R is a Vaccinia virus encoded decoy protein that binds type I IFNs and antagonises their function^17,38^. B18R treatment of the XRN1 KO cells reduced expression of RIG-I, MDA5 and downstream factors, including STAT1 and pSTAT1 to similar levels as WT cells (**Fig.4A**). Consistent with these observations B18R significantly reduced gene expression of the well-known ISGs, MxA and RSAD2 (**Fig.4A**). To assess the impact of secreted IFNs on HDV replication we treated the infected cells with B18R and measured viral RNAs at 6 dpi. B18R had a minimal effect of HDV replication in WT cells, in contrast we noted a significant 2- or 4-fold increase in viral RNAs in the KO22 and KO35 cells, respectively (**Fig.4B**). These data indicate that the secreted IFNs supress HDV infection of XRN1 depleted hepatoma cells.

**Figure. 4.**
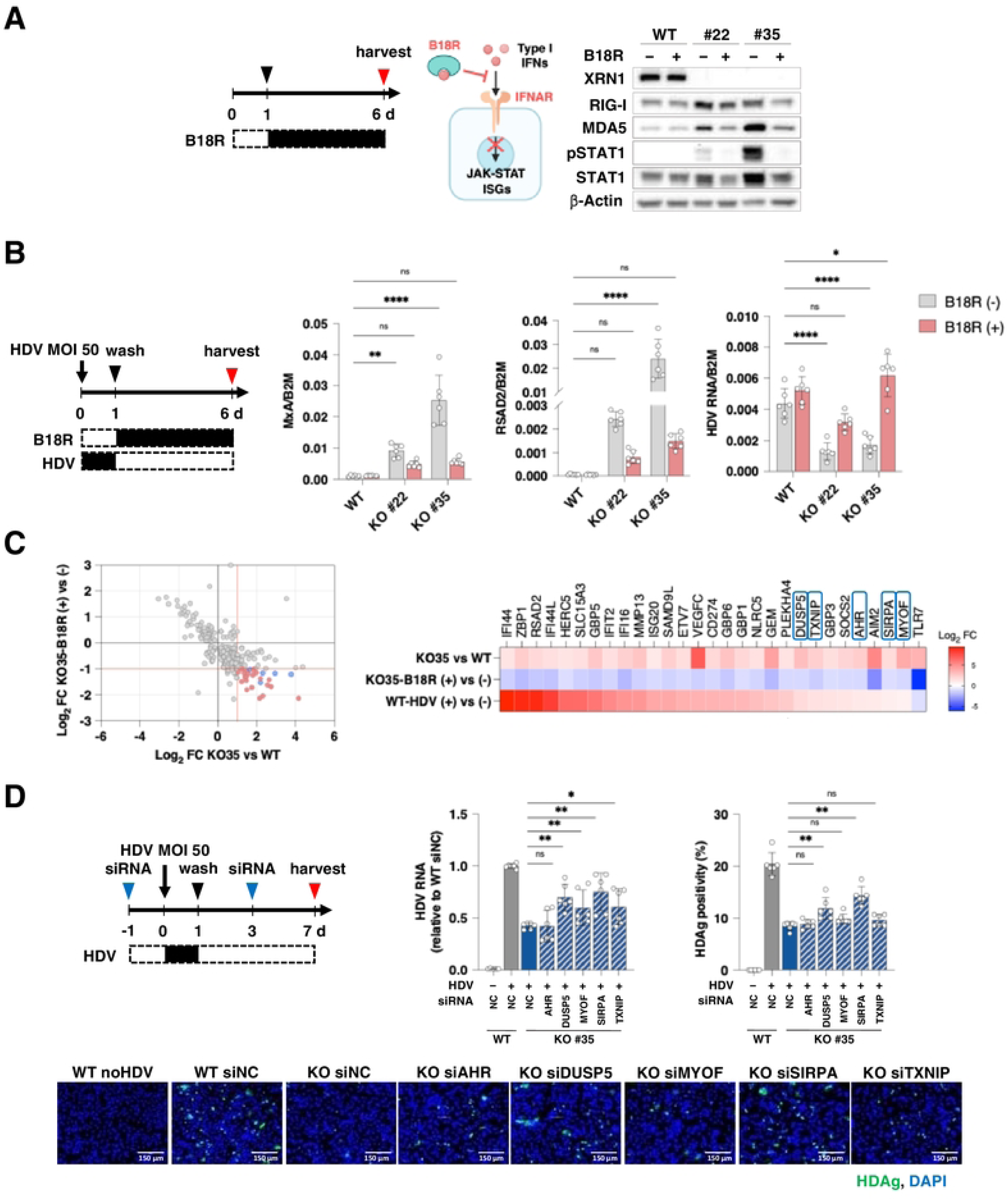
DUSP5 and SIRPA induced by the dsRNA sensing pathway restrict HDV replication. (**A**) **Effect of B18R on the dsRNA pathway.** Differentiated HepG2-NTCP WT or XRN1 KO cells were incubated in the presence or absence of 10 ng/mL of B18R for 5 days. Protein levels of XRN1, RIG-I, MDA5, pSTAT1, STAT1 and β-Actin were assessed by western blotting. (**B**) **Effect of B18R on HDV replication.** Differentiated HepG2-NTCP WT or XRN1 KO cells were infected with HDV (MOI = 50) for 24 h, then incubated in the presence or absence of 10 ng/mL of B18R for 5 days. RNA levels of HDV and the IFN pathway factors MxA and RSAD2 were assessed by RT-qPCR. The data were normalized to the housekeeping gene, *B2M* (mean ± SD, n = 6 from 3 independent experiments, Mann-Whitney tests with Bonferroni correction * *p* < 0.05, ** *p* < 0.01, \*\*\**p* < 0.005, **** *p* < 0.0001). (**C**) **Differentially expressing ISGs.** The samples shown in Figure 3C (WT and KO #35) were also analyzed by RNAseq. The log_2_FC of ISGs in XRN1 KO #35 cells relative to WT (X axis), was compared to the log_2_FC observed between B18R treated and untreated XRN1 KO #35 cells (Y axis). The factors identified by red and blue points (X axis >1 and Y axis <-1, marked by dashed lines) are shown in the heatmap. (**D**) **siRNA screening of selected ISGs.** HepG2-NTCP WT and KO #35 cells were differentiated for 2 days and then transfected with a non-targeting control siRNA (NC) or siRNAs (AHR, DUSP5, MYOF, SIRPA and TXNIP, 50nM) for 6h. After 24 h cells were infected with HDV (MOI = 50) for 24 h, cells washed and incubated for 6 dpi. siRNAs were re-transfected at 3dpi. HDV RNA and HDAg were detected by RT-qPCR and immunofluorescence staining, respectively. The RT-qPCR data were normalized to a housekeeping gene, *B2M* (mean ± SD, n = 6 from 3 independent experiments, Mann-Whitney tests with Bonferroni correction for multiple comparisons. * *p* < 0.05, ** *p* < 0.01). The images show HDAg (Green) and nuclei (Blue) detection by immunofluorescence staining and are from one of 3 independent experiments.

### Identification of anti-viral ISGs

Several reports have shown that HDV infection activates IFN-β and IFN-λ expression and induces a broad spectrum of ISGs with limited impact on viral replication^23,39^. Reassuringly, the minimal effect of B18R on HDV infection of WT cells supports this conclusion (**Fig.4B**). Although Peg-IFN-α is used to treat chronic hepatitis D, IFN-α or IFN-λ treatment had a minimal impact on HDV replication in experimental model systems^23^. Since the dsRNA sensing pathway induced by XRN1 depletion repressed HDV replication, we reasoned that analysing the transcriptome of B18R-treated HepG2-NTCP cells could identify ISGs with anti-viral activity. We identified 23 upregulated and 13 down regulated ISGs in XRN1 KO35 cells, this phenotype was lost following B18R treatment (**Fig.4C**). Excluding ISGs that were also induced by HDV identified five XRN1-specific genes whose influence on HDV replication in KO22 and KO35 cells were assessed. Aryl hydrocarbon receptor (*AHR*), dual Specificity Phosphatase 5 (*DUSP5*), myoferlin (*MYOF*), signal regulatory protein Alpha (*SIRPA*), and thioredoxin interacting protein (*TXNIP*) were siRNA silenced prior to infection and confirmed by RT-qPCR (**Supplementary Fig. 1A**). Infection was assessed by measuring HDV RNA and HDAg expression using an antibody that recognises both L- and S-HDAg (**Fig.3B**). HDV replication was reduced in both KO22 and KO35 cells compared with WT cells following delivery of a non-targeting control siRNA (NC), consistent with our earlier observations (**Fig.4D, Supplementary Fig. 1B**). In contrast, silencing MYOF or TXNIP increased HDV replication in the KO35 cells, but these effects did not reach significance in the KO22 cells. Silencing DUSP5 and SIRPA significantly increased both HDV RNA and HDAg expression in the KO lines (**Fig.4D, Supplementary Fig. 1B**), confirming their role in the regulation of HDV infection. In conclusion we have identified a dual mechanism for XRN1 regulation of HDV infection. Firstly, via modulating ADAR1-p150 expression and secondly acting as a rheostat to maintain steady state dsRNA levels and associated antiviral host innate immune responses.

## DISCUSSION

XRN1 is the major cytoplasmic 5′-3′ exoribonuclease responsible for maintaining RNA homeostasis through degradation of cellular transcripts. Recent studies identified a role for XRN1 in the regulation of endogenous retroviral transcripts that can activate dsRNA sensing pathways^26,27^. Here, we expand these studies to focus on XRN1 as a regulator of innate immune restriction of HDV. Loss of XRN1 triggered endogenous dsRNA accumulation, activation of dsRNA sensing pathways, and the induction of an antiviral state that restricts HDV replication. These findings demonstrate a role for endogenous RNA metabolism in defining the outcome of HDV infection.

XRN1 has been reported to exert both antiviral and proviral functions during RNA virus infection, highlighting its context-dependent role in host–virus interactions. XRN1 directly degrades the genomic RNAs of several mammalian RNA viruses, including Newcastle disease virus, encephalomyocarditis virus, Sendai virus, and Coxsackievirus B, thereby restricting viral replication^18–21^. Consistent with the antiviral pressure imposed by XRN1, several viruses have evolved mechanisms to evade its activity. For example, poliovirus promotes proteasomal degradation of XRN1, whereas flaviviruses generate XRN1-resistant subgenomic RNAs (xrRNAs) through structured elements in their 3′ untranslated regions that stall XRN1 and attenuate host antiviral responses^40–42^.

XRN1 can promote viral infection by limiting innate immune activation. In measles virus infected cells, XRN1 is recruited to cytoplasmic viral inclusion bodies, where it prevents the accumulation of immunostimulatory dsRNA and subsequent PKR activation^43^. Depletion of XRN1 during vaccinia virus infection enhanced dsRNA accumulation, leading to activation of PKR and RNase L and suppression of viral replication through translation arrest and RNA degradation^16^. XRN1 has been reported to interact with the IAV non-structural (NS) protein and to enhance its suppression of RIG-I activation and IFN-β production in lung A549 cells^44^. In contrast, we observed a modest increase in IAV replication in XRN1 KO HeG2-NTCP cells (**Fig.2C**), suggesting that the consequences of XRN1 depletion for viral replication are dependent on the cellular context.

XRN1 depletion has been reported to activate diverse innate immune pathways in a range of cell types. In the colorectal cancer cell line NCI-H747, depletion of MAVS or PKR but not RNase L partially rescued cell viability caused by XRN1KO, implicating MAVS and PKR pathways^26^. PKR was identified as the predominant mediator of cell death following XRN1 deletion in the non-small cell lung cancer cell lines NCI-H1650 and HCC366^27^. In contrast, in IAV infected lung A549 cells RIG-I was activated following XRN1 depletion^44^. We noted increased expression of phosphorylated PKR in the KO HepG2 cells (**Fig.4**) consistent with its activation, IFN expression and elevated RIG-I and MDA5 levels. Collectively, these observations illustrate a homeostatic role for XRN1 in the regulation of dsRNA levels and down stream IFN signaling.

One potential determinant of the context-dependent responses is the abundance and composition of endogenous dsRNAs. Previous studies have shown that basal dsRNA levels vary among cancer cell lines and this associates with PKR-associated lethality following IFN treatment^26,27^. Moreover, increasing endogenous dsRNA through epigenetic perturbation, including treatment with the DNA-demethylating agents 5-aza-2’-deoxycytidine (5-AZA-CdR) or cyclin-dependent kinase 4 and 6 (CDK4/6) inhibitors, alters cellular sensitivity to XRN1 depletion^26^. These observations suggest that both the abundance and molecular origin of endogenous dsRNA may determine which innate immune sensors are engaged following loss of XRN1. Characterizing transposable element-derived transcripts together with sensor-specific dsRNA populations across different cellular contexts will be important for understanding XRN1-dependent RNA homeostasis. Comparison of our data with previously reported XRN1-dependent models is limited by the differences in experimental systems used and the limited availability of transcriptomic datasets. Future comparative analyses across cell types will help to define the determinants that specify the downstream response to XRN1 depletion.

ADAR1-mediated adenosine-to-inosine (A-to-I) RNA editing destabilizes dsRNA structures and prevents aberrant activation of innate immune sensors, thereby functioning as a major suppressor of viral mimicry^26,27,33,45–47^. Recent genetic studies have revealed a close functional relationship between XRN1 and ADAR1 in regulating endogenous dsRNA and innate immunity^26,27^. We found that XRN1 ablation induced ADAR1 expression and increased A-to-I editing of both viral and host RNAs. We speculate that enhanced ADAR1 activity may constitute the compensatory feedback mechanism that counteracts excessive dsRNA accumulation and innate immune activation following disruption of XRN1-mediated RNA decay. Thus, XRN1-mediated RNA degradation and ADAR1-mediated RNA editing may represent complementary mechanisms for maintaining cellular dsRNA homeostasis. Further studies will be required to determine how these pathways are coordinated and how their interplay influences antiviral immunity during HDV infection.

To date, several ISGs have been reported to facilitate rather than restrict HDV replication. ADAR1 edits the HDAg stop codon to generate the large HDAg (L-HDAg) that is essential for virion assembly and the ISG moloney leukaemia virus 10 protein (MOV10) promotes HDV RNA synthesis through its association with HDAg (reviewed in^48^). More recently, Janus kinase 1 (JAK1), a central component of cytokine signalling, was reported to facilitate HDV replication through activation of ERK1/2 and phosphorylation of small HDAg (S-HDAg)^49^. Our transcriptomic profiling, combined with functional siRNA screening, identified DUSP5 and SIRPA as previously unrecognized ISGs that restrict HDV replication. DUSP5 is a nuclear phosphatase that dephosphorylates and inactivates ERK1/2, raising the possibility that it restricts HDV by counteracting the proviral JAK1–ERK1/2 signalling axis^49^. DUSP5 was reported to inhibit dengue and Zika virus infection by negatively regulating the ERK-MLCK-Myosin IIB signalling axis and suppression of F-actin rearrangements^50^. SIRPA is a cell surface receptor that inhibits macrophage phagocytic activity and has been reported to restrict Zika virus, Vesicular stomatitis virus and Lymphocytic choriomeningitis virus internalization^51,52^. We previously reported that XRN1 depletion had a minimal effect on HBV entry^28^ and since HDV shares the same NTCP-dependent internalization pathway, SIRPA is likely to restrict HDV through a post-entry mechanism. Defining the molecular mechanisms by which DUSP5 and SIRPA restrict HDV is an important direction for future studies.

Collectively, our study identifies XRN1-dependent RNA homeostasis as a determinant of innate immune control of HDV. Disruption of XRN1-mediated RNA decay induces dsRNA accumulation and IFN signaling pathways that establish an antiviral state and limit HDV replication via DUSP5 and SIRPA dependent pathways. Selective activation or therapeutic exploitation of such antiviral pathways, rather than broad IFN stimulation, may represent an alternative strategy for strengthening host-mediated control of chronic hepatitis D.

### Limitation of the study

One limitation is that our transcriptomic analysis identified transposable element transcripts induced following XRN1 depletion but does not define the immunogenic dsRNAs responsible for activating individual RNA sensors. Identification of sensor-bound RNAs by biochemical approaches, such as MDA5 immunocapture followed by RNA sequencing^32^, together with analogous analyses for PKR and OAS/RNase L, would be required to identify the dsRNA species that drive antiviral signaling in hepatocytes.

## MATERIALS AND METHODS

### Reagents

Recombinant B18R protein and Poly(I:C) were purchased from InvivoGen, Human IFN-α2a, Human IFN-β1a and Human IFN-γ1b from Miltenyi Biotec. The antibodies along with their supplier are: XRN1 (Thermo Fisher Scientific), HDAg (Kerafast), ADAR1 (Santa Cruz), RIG-I (AdipoGen), MDA5 (Caltag Medsystems), MAVS (Cell Signaling), STAT1 (Cell Signaling), phospho-STAT1 (Tyr701, Cell Signaling), dsRNA (rJ2, Merck), β-Actin (Sigma), HBsAg (Abcam), anti-mouse Immunoglobulin HRP (Dako), anti-rabbit IgG HRP (Cytiva), anti-mouse IgG HRP (Abcam), anti-rabbit IgG (Alexa Fluor 488, 594 and 633, Thermo Fisher Scientific). The XRN1 expression plasmid (pLNHA-C1-HsXRN1, #66596) and HDV genome plasmid (pSVLD3, #29335) were purchased from Addgene (plasmid). Silencing RNAs were purchased from Thermo Fisher Scientific.

### Cell lines

HepG2-NTCP XRN1-WT and KO cells, HuH-7 cells, HEK293-ISRE-Luc reporter and HEK293-P125 (IFNβ-Luc promoter) cells were maintained in Dulbecco’s Modified Eagles Medium (DMEM) containing Glutamax supplemented with 10% Fetal Bovine Serum, 50 U/ml Penicillin/Streptomycin, and non-essential amino acids (all reagents from Thermo Fisher Scientific). Cells were maintained in 5% CO_2_ and 18% O_2_ at 37°C.

### Transfection

HEK293T cells were transfected with siRNAs using DharmaFECT 4 Transfection Reagent (Horizon Discovery). HepG2-NTCP cells were transfected with plasmids using either polyethylenimine (PEI) or FuGENE HD Transfection Reagent (Promega), according to the manufacturer’s protocol.

### Luciferase assay

Cells were washed with PBS and lysed using Passive Lysis Buffer (Promega). Luciferase activity was determined using a Luciferase Assay System (Promega), according to the manufacturer’s protocol.

### Detection of IFN activity in cell culture supernatant

HepG2-NTCP WT and KO cells were seeded and cultured in a 24 well plate for 72h. HEK293T cells stably expressing an IFN-stimulated response element (ISRE)-derived reporter were seeded in a 96 well plate and treated with IFNα or with the supernatants from HepG2-NTCP XRN1 WT or KO cells for 48h to detect luciferase activity.

### Preparation of HDV

Twenty-four hours prior to transfection, Huh-7 cells were seeded in complete medium at a density of 6E+6 cells/T75 flask. The next day, cells were transfected with 5 μg of pSVLD3 and 5 μg of pT7HB2.7 plasmids using 30 μL of FuGENE-HD reagent (Promega). Following overnight incubation, cells were washed twice with PBS and the medium replaced with 12.5 mL of complete medium. Supernatant was collected at 3, 6, 9 and 12 days post-transfection and concentrated to 1/50^th^ of the original supernatant volume using 50 kDa ultra-centrifugal filters (Merck) and used as HDV inoculum. To quantify genome copy number, RNA from 3 μL of concentrated HDV inoculum was extracted using a Viral RNA Mini Kit (QIAGEN), and reverse transcribed using a cDNA synthesis kit (PCR Biosystems), according to the manufacturer’s protocol (25°C, 10 min; 42°C, 15 min; 48°C, 15 min; 85 °C, 10 min) and quantified using a SyGreen Blue Mix (PCR Biosystems) with a program of 95°C, 2 min; 45 cycles of 95°C, 5 sec; 60°C, 30 sec, using primers specific for HDV (5’-TCTTCCTCGGTCAACCTCTT-3’ and 5’-ACAAGGAGAGGCAGGATCAC-3’). Genome copy number was enumerated against a standard curve of pSVLD3 plasmid of known concentration.

### HDV infection

HepG2-NTCP cells were seeded on collagen-coated plasticware and cultured in DMEM containing 2.5% DMSO for 72h. Cells were inoculated with HDV (MOI of 50 genome copies) in the presence of 4% PEG8000 and 2.5% DMSO for 24 h. Viral inoculum was removed and cells washed three times with PBS. Infected cells were maintained in DMEM containing 2.5% DMSO in 5% CO_2_ and 18% O_2_ at 37°C, with subsequent treatments and experimental timelines as described.

### IAV, SARS-CoV-2, HCoV and EMCV infection

Viral stocks were obtained from BEI Resources, NIAID, NIH. HepG2-NTCP WT or KO22 cells were seeded on collagen coated plasticware and infected with Influenza A virus A (IAV) PR8-GFP, SARS-CoV-2 (VIC-01 mNeonGreen), HCoV-OC43 or Encephalomyocarditis virus (EMCV) at an MOI of 1. For PR8-GFP and SARS-CoV-2 infections, cells were fixed in 4% PFA 24h post infection and GFP fluorescence quantified by CLARIOstar with data normalised to a DAPI counter stain. For OC43 and EMCV viral titres were quantified by plaque assay on Vero as previously described^53,54^.

### Cytotoxicity

Cytotoxicity was determined using a Lactate dehydrogenase CytoTox 96 Non-Radioactive Cytotoxicity Assay (Promega), according to the manufacturer’s protocol.

### Immunofluorescence staining

Cells were washed twice with PBS and fixed with 4% formaldehyde in PBS for 30 min. After fixation, samples were washed three times with PBS, permeabilized with 0.25% Triton X-100 in PBS for 20 min and blocked in 3% bovine serum albumin (BSA) in PBS for 30 min at room temperature (R.T.). Primary antibodies were diluted in 1% BSA in 0.1% Tween 20 containing PBS (PBS-T) and incubated with the cells at 4°C overnight or at R.T. for 1 h. Cells were washed three times with PBS and two times with PBS-T. Secondary antibodies and DAPI were diluted to 1:500 and 1:5000 respectively in 1% BSA/PBS-T. Cells were incubated with the antibodies at R.T. for 1 h in the dark and washed three times with PBS, two times with PBS-T, and finally two additional times with PBS. Coverslips were mounted on to glass slides with VECTASHIELD Antifade Mounting Medium (Thermo Fisher Scientific). Images were acquired using EVOS M5000 Cell Imaging Systems (Thermo Fisher Scientific) or SpinSR SoRa cell imaging systems (Olympus) and processed in ImageJ.

### Quantitative cDNA PCR (RT-qPCR)

Total cellular RNA was extracted using an RNeasy kit (Qiagen) according to the manufacturer’s instructions. RNA from plasmid-transfected cells was treated with TURBO DNase-free (Thermo Fisher Scientific) according to the manufacturer’s instructions. RNA was reverse transcribed using a cDNA synthesis kit (PCR Biosystems) using a program of 25°C, 10 min; 42° C, 15 min; 48°C, 15 min; 85°C, 10 min. Gene expression was quantified using a SyGreen Blue Mix (PCR Biosystems) using a qPCR program of 95°C, 2 min; 45 cycles of 95°C, 5 sec; 60°C, 30 sec. Oligonucleotides are listed in Supplementary Table 2. Changes in gene expression were calculated relative to the housekeeper gene, *β 2-microglobulin* (ΔCt method).

### Single molecule in situ hybridization (smiFISH)

SmiFISH was performed as previously described^37^. Cells were seeded on coverslips and infected with HDV. At 6dpi, coverslips were washed twice with PBS and fixed with 4% formaldehyde in PBS for 30 min. After fixation, coverslips were washed three times with PBS, permeabilized with 0.25% Triton X-100 in PBS for 20 min at R.T, treated with RNAsecure (Thermo Fisher Scientific) for 30 min at RT, and pretreated with a FISH wash buffer (10% formamide, 2X SSC (Invitrogen)). A Cy3 conjugated FLAP probe was annealed to the smiFISH probe mix in 1x NEB 3 buffer (New England Biolabs) in a thermocycler at 85°C, for 3 min, 65°C, for 3 min and 25°C, for 5 min. Coverslips were hybridized with annealed probes overnight in a buffer containing 10% formamide, 2X SSC, 8% dextran sulfate, 0.34mg/mL tRNA (Merck), 1mM Ribonucleoside Vanadyl Complex (New England Biolabs), 0.01% BSA. After hybridization, coverslips were washed in FISH wash buffer twice for 15 min at 37°C, then subsequently washed for 15 min three times in a buffer containing 2x SSC 0.1% Tween 20. Coverslips were treated with primary antibody overnight, then with secondary antibody and DAPI for 1h.

### SDS-PAGE and western blotting

Samples were lysed in RIPA buffer (50 mM Tris (pH 8.0), 150 mM NaCl, 1% Nonidet P-40 (NP-40), 0.5% sodium deoxycholate, and 0.1% sodium dodecyl sulphate supplemented with cOmplete Protease Inhibitor Cocktail (Merck). 4x Laemmli reducing buffer was added to samples before heating at 95°C for 10 min. Proteins were separated on 8% polyacrylamide gels and transferred to activated 0.45 µm PVDF membranes (Amersham). Membranes were blocked in 5% skimmed milk and proteins detected using specific primary and HRP-conjugated secondary antibodies. Signals were realised using a SuperSignal West Pico chemiluminescent substrate kit (Pierce) and images acquired on a G:Box mini (Syngene).

### Poly-A RNA sequencing and analysis

HepG2-NTCP cells were differentiated by treating with 2.5% DMSO for 3 days and infected with mock or HDV (MOI = 50) for 24h. After infection, the cells were washed and incubated in the presence or absence of 10 ng B18R for 5 days. RNA was extracted using an RNeasy kit with on-column DNase treatment in accordance with the manufacturer’s instructions (Qiagen). mRNA libraries were prepared by poly A enrichment and analysed by 300bp paired read sequencing on a NovaSeq X Plus Sequencer (Novogene). Sequences were aligned using minimap2^55^ to a referent cDNA dataset consisting of a human transcriptome, Homo sapiens, assembly GRCh38 (file: Homo_sapiens.GRCh38.cdna.abinitio.fa.gz), HBV (NC_003977.2) and HDV (NC_001653.2), together with the sequence of a plasmid pUC57 (Y14837) as a contamination control. For HDV the number of AUC (parental) and ACC (edited) codons in the alignment were determined. For the host transcriptome the relative frequencies of ADAR1 editing were measured using a database of known ADAR1 targets^56^. The levels of expression of transposable elements in the WT and KO transcriptomes were determined and compared using the TEcounts and TEtranscripts packages respectively^57^. The RNA-seq data is available as an SRA at NCBI (BioProject ID: PRJNA1228332).

### Total RNA sequencing and analysis

HepG2-NTCP WT or KO 22 cells were seeded on collagen coated plasticware and cultured in DMEM containing 2.5% DMSO for 6 days. Cells were harvested and RNA extracted using an RNeasy kit with on-column DNase treatment in accordance with the manufacturer’s instructions (Qiagen). Total RNA libraries were prepared by ribosomal depletion, cDNA synthesised using random hexamers, adapter tailed and size selected (Novogene). Sequences were obtained by 300bp paired read sequencing on a NovaSeq X Plus Sequencer. Sequencing reads were processed using a modified fork of nf-core/rnaseq (v3.18.0^58,59^). Adapters and homopolymer tails were trimmed, and reads were quality-filtered using fastp (v0.23.4^60^). Read quality was assessed using FastQC (v0.12.1). Following rRNA filtering, reads were aligned using STAR (v2.7.11b^61^) to the CHM13v2.0 assembly^62^, using GENCODE annotation (v49^63^) transferred from GRCh38 with Liftoff (v1.6.3^64^). To characterize A-to-I editing, a shared editome was first constructed from observed A-to-G mismatches. Sites with coverage ≥ 8 and editing frequencies of 5-45% that were detected in at least three samples or previously reported in REDIportal^65^ were clustered using DBSCAN (scikit-learn v1.8.0; Pedregosa et al., 2011; ε = 120 bp). Differential editing between parental and XRN1-KO cells was then assessed using DEXSeq (v1.56.0^66^). Testing required at least two replicates per condition with mean coverage ≥ 8 per editing cluster. Clusters were then classified as significant (Benjamini-Hochberg FDR ≤ 0.05, median editing-frequency difference ≥ 5%) or as condition-specific if they met the editing-frequency and coverage thresholds in one condition but not the coverage threshold in the other. The total RNA-seq data is available as an SRA at NCBI (BioProject ID: PRJNA1539819).

## FUNDERS

Research in the McKeating laboratory is funded by the Wellcome Trust (Investigator Award 200838/Z/16/Z and Discovery Award 225198/Z/22/Z), MRC project grant (UKRI3656) and the Chinese Academy of Medical Sciences (CAMS) Innovation Fund for Medical Science, China (grant no. 2024-I2M-2-001-1). MB is funded by a Kings College Hospital-Oxford University research agreement. XZ is funded by a CAMS training fellowship. Research in the Rehwinkel laboratory is funded by the UK Medical Research Council (MR/Y013212/1). Research in the Wing laboratory is funded by BBSRC (UKRI3565) and the Chinese Academy of Medical Sciences (CAMS) Innovation Fund for Medical Science, China (grant no. 2024-I2M-2-001-1). AF receives funding from the Clarendon Fund, the Radcliffe Department of Medicine, and Keble College.

## ACKNOWLEDGEMENTS

We would like to thank: Stephan Urban (University of Heidelberg) for HepG2-NTCP cells; Pierre Khalfi and Karim Majzoub (University of Montpellier) for smiFISH probes and protocol; Alfredo Castello (Centre for Virus Research, University of Glasgow) for XRN1 Cas9 KO reagents; Camille Sureau (University of Tours) for pT7HB2.7 plasmid and Rebecca Berrens (University of Oxford) for advice on TE mapping.

## AUTHOR CONTRIBUTIONS

S.T. designed and conducted experiments and co-wrote the MS; M.B. provided technical support; P.A.C.W. designed and conducted experiments; A.F, X.Z and P.B. analyzed data; J.R. advised on experimental design and provided reagents; J.A.M. designed study and co-wrote MS.

---

**Supplementary Figure 1. siRNA screening of selected ISGs in XRN1 KO22 cells.** (**A**) HepG2-NTCP WT, KO #22 and KO #35 cells were differentiated for 2 days and transfected with non-targeting control siRNA (NC) or siRNAs (AHR, DUSP5, MYOF, SIRPA and TXNIP, 50nM) for 6h. siRNAs were transfected again after 3 days. After 6 days RNA was extracted and levels of AHR, DUSP5, MYOF, SIRPA and TXNIP quantified by RT-qPCR with normalization to the housekeeping gene, *B2M* (mean ± SD, n = 6 from 3 independent experiments). (**B**) **siRNA screening of selected ISGs in KO #22 cells.** HepG2-NTCP WT and KO #22 cells were differentiated for 2 days and then transfected with a non-targeting control siRNA (NC) or siRNAs (AHR, DUSP5, MYOF, SIRPA and TXNIP, 50nM) for 6h. After 24 h cells were infected with HDV (MOI = 50) for 24 h, washed and incubated for 6 dpi. siRNAs were re-transfected at 3dpi. HDV RNA and HDAg were detected by RT-qPCR and immunofluorescence staining, respectively. The RT-qPCR data were normalized to a housekeeping gene, B2M (mean ± SD, n = 6 from 3 independent experiments, Mann-Whitney tests with Bonferroni correction for multiple comparisons, ** p< 0.01). The images show HDAg (Green) and nuclei (Blue) detection by immunofluorescence staining and are from one of 3 independent experiments.

**Supplementary Table 1. Genes listed in the IFN-α and -γ pathway responses by GSEA.**

**Supplementary Table.2 RT-qPCR primer sequences.**

## Notes

### Competing Interest Statement

The authors have declared no competing interest.

